# Improved pangenomic classification accuracy with chain statistics

**DOI:** 10.1101/2024.10.29.620953

**Authors:** Nathaniel K. Brown, Vikram S. Shivakumar, Ben Langmead

**Affiliations:** Department of Computer Science, Johns Hopkins University, Baltimore MD 21218

**Keywords:** Pangenomics, text indexing, sequence classification

## Abstract

Compressed full-text indexes enable efficient sequence classification against a pangenome or tree-of-life index. Past work on compressed-index classification used matching statistics or pseudo-matching lengths to capture the fine-grained co-linearity of exact matches. But these fail to capture coarse-grained information about whether seeds appear co-linearly in the reference. We present a novel approach that additionally obtains coarse-grained co-linearity (“chain”) statistics. We do this without using a chaining algorithm, which would require superlinear time in the number of matches. We start with a collection of strings, avoiding the multiple-alignment step required by graph approaches. We rapidly compute multi-maximal unique matches (multi-MUMs) and identify BWT sub-runs that correspond to these multi-MUMs. From these, we select those that can be “tunneled,” and mark these with the corresponding multi-MUM identifiers. This yields an ℴ(*r* + *n/d*)-space index for a collection of *d* sequences having a length-*n* BWT consisting of *r* maximal equal-character runs. Using the index, we simultaneously compute fine-grained matching statistics and coarse-grained chain statistics in linear time with respect to query length. We found that this substantially improves classification accuracy compared to past compressed-indexing approaches and reaches the same level of accuracy as less efficient alignmentbased methods.

## 1 Introduction

Read classification is at the core of sequencing data analyses like taxonomic classification, host sequence depletion, and nanopore adaptive sampling. Meanwhile, databases of reference sequences are growing thanks to improvements in long-read sequencing [13, 16]. This has spurred advances in compressed indexing methods that specialize in repetitive collections, such as intra-species pangenomes or inter-species collections from across the tree of life. Previous methods like SPUMONI and SPUMONI 2 [2, 1] support efficient binary and multi-class classification of reads against compressed indexes. These build on the *r*-index [12, 14], a compressed index that grows with the amount of distinct sequence in the reference collection. They also build on the MONI algorithm [23], which enables computation of matching statistics (MSs). The SPUMONI study suggested an alternative to matching statistics called pseudo-matching lengths (PMLs), which are an approximate version of MSs that can be computed more efficiently.

While these methods have had success, they are currently less accurate than alignment-based classification methods. For example SPUMONI 2’s accuracy trailed that of a minimap2-alignment-based method [15] by about 2.5 percentage points on sensitivity and about 0.8 percentage points on specificity for both microbial mock-community and human microbiome datasets. We hypothesized that this gap is due to the alignment-based tools’ facilities for *chaining* matches (i.e. seed hits) into sets of matches that are colinear with respect to the reference. While this can improve accuracy, it is also expensive. Chaining requires *O*(*m* log *m*) time with respect to the number of seeds *m*, with some practical methods like minimap2 employing a simpler but theoretically slower *O*(*m*^2^)-time algorithm. Studies have shown that for especially long reads, chaining is the bottleneck of minimap2 [24].

To date, compressed-indexing-based methods have lacked an equivalent chaining facility. By computing PMLs rather than full matching statistics, and by excluding the suffix-array sample from the index, SPUMONI foregoes any knowledge of match location with respect to the reference. Furthermore, in a full-text pangenome context, “location” information becomes complex, as seed hits to homologous regions in different haplotypes would present as distant matches index without a unifying coordinate system. This further taxes the computation required to perform chaining.

Sigmoni [26] introduced a method for approximate location-based match clustering by dividing documents into non-overlapping partitions (“shreds”) and augmenting the index with a shred-level document array specifying each suffix’s shred of origin. While this method improves classification for single reference genomes, it does not extend to pangenomic references as related genomes can differ drastically either in their lengths and/or in their structural-variant content, affecting the linear coordinate system and misaligning homologous shreds. Further, since the shreds have preset widths that are not a function of genome length, the shred-level document array requires *O*(*r* log *n*) space, where *n* is the total collection length.

We present a novel method and structure called the col-BWT that enables compressed indexes to generate both matching statistics and chain statistics simultaneously and in linear time with respect to the query length. Chain statistics complement the fine-grained co-linearity information inherent in MSs and PMLs by additionally conveying whether matches are co-linear with respect to the reference sequences in the index. Further, the col-BWT fits in *O*(*r* + *n/d*) space, where *d* is the number of indexed sequences, *n* is the total length of the indexed sequences, and *r* is the number of runs in the collection’s BWT. Past work shows that growth proportional to *r* allows for scaling to large pangenomes; the SPUMONI 2 study, for example, showed that an index of 10 human haplotypes fit in 10.2 GB. The additive *n/d* factor equals the average length of the sequences in the collection, which does not grow substantially with the collection size.

Besides these theoretical advances, we show experimentally that our col-BWT index fits in 46.3 GB when built over 10 human haplotype sequences and 48.28 Gb when built over 32 human haplotype sequences from the HPRC [16]. Further, we show that use of chain statistics allows col-BWT to achieve classification accuracy that substantially closes the gap with an alignment-based method based on minimap2. Specifically, col-BWT achieves a smaller 0.6 percentage-point deficit on sensitivity, and achieves slightly superior specificity when classifying with respect to a pangenome consisting of 10 HPRC haplotypes.

## 2 Preliminaries

### 2.1 Notation

An array *A* of | *A*| = *n* elements is represented as *A*[1..*n*]. {*A*} is the set derived from *A* of size |{*A*}| . For *x* ∈ℕ, let [1, *x*] represent the array [1, 2, … , *x*]. A *string S*[1..*n*] is an array of symbols drawn from an ordered alphabet *Σ* of size *σ*. We use *S*[*i*..*j*] to represent a *substring S*[*i*] … *S*[*j*]. For strings *S*_1_, *S*_2_ let *S*_1_*S*_2_ represent their concatenation. A *prefix* of *S* is some substring *S*[1..*j*], where a *suffix* is some substring *S*[*i*..*n*]. The *longest common prefix* (*lcp*) of *S*_1_,*S*_2_ is defined as *lcp*(*S*_1_, *S*_2_) = max{*j* | *S*_1_[1..*j*] = *S*_2_[1..*j*]}. A *text T* [1..*n*] is assumed to be terminated by the special symbol $ ∉ *Σ* of least order so that suffix comparisons are well defined. We use the RAM word model, assuming machines words of size 𝒲= *Θ*(log *n*) with basic arithmetic and logical bit operations in *O*(1)-time.

### 2.2 Suffix Array and Burrows-Wheeler Transform

The *suffix array* (SA) [17] of *T* [1..*n*] is a permutation SA[1..*n*] of [1, *n*] such that SA[*i*] is the starting position of the *i*th lexicographically smallest suffix of *T* . The *longest common prefix array* (LCP) stores the *lcp* between lexicographically adjacent suffixes in *T* [1..*n*], i.e., LCP[1..*n*] such that LCP[1] = 0 and LCP[*i*] = *lcp*(*T* [SA[*i*− 1..*n*], *T* [SA[*i*..*n*]) for *i >* 1. We use the term *document* to refer to a single genome from among the many being indexed. Let a collection of documents 𝒟= {*T*_1_, *T*_2_, … , *T*_*d*_} be represented by their concatenation^1^ *T* [1..*n*] = *T*_1_*T*_2_ … *T*_*d*_$. The *document array* (DA) [19] is defined as DA[1..*n*] where DA[*i*] stores which document *T* [SA[*i*]..*n*] begins in. We use *d* to refer to the number of sequences/documents in a collection.

The *Burrows-Wheeler Transform* (BWT) [9] is a permutation BWT[1..*n*] of *T* [1..*n*] such that BWT[*i*] = *T* [SA[*i*] − 1] if SA[*i*] *>* 1, otherwise BWT[*i*] = *T* [*n*] = $. The BWT is reversible by using the *last-to-first* (LF) mapping LF(*i*), a permutation over [1, *n*] satisfying SA[LF(*i*)] = (SA[*i*] − 1) mod *n*. The *first-to-last* (FL) mapping is its inverse such that FL(*i*) satisfies SA[FL(*i*)] = (SA[*i*] + 1) mod *n*. Performing successive LF steps is referred to as *backwards stepping* where we use the recursive notation LF^*x*^(*i*) = LF^*x*−1^(LF(*i*)) for *x >* 0 and LF^0^(*i*) = *i. Forward stepping* is defined symmetrically using FL. These permutations allow substrings of the original text to be extracted.

### 2.3 *r*-index and Move Structure

Let *r* be the number of maximal equal character runs of the BWT. The *run-length encoded BWT* (RLBWT) is an array RLBWT[1..*r*] of tuples where RLBWT[*i*].*c* is the character of the *i*th BWT run and RLBWT.*ℓ* its length. In the *r*-index study by Gagie et al. [12], the RLBWT is combined with run-sampled SA values to answer substring queries, specifically locate queries, efficiently in *O*(*r*) words of space. It supports LF in *O*(log log _𝒲_ (*n/r*)-time. Nishimoto and Tabei improved this to optimal-time^2^ substring queries in *O*(*r*)-space using their *move structure* [20]. For an arbitrary permutation *π* on [1, *n*], where *b* is the number of positions *i∈* [1, *n*] such that either *i* = 1 or *π*(*i* −1) ≠ *π*(*i*) 1 mod *n*, the move structure computes *π* in *O*(1)-time and *O*(*b*)-space. For LF and FL these positions correspond directly to run boundaries; they can computed in *O*(1)-time and *O*(*r*)-space.

### 2.4 Fine-Grained Co-linearity Statistics

The *matching statistics* (MS) of a pattern *P* [1..*m*] with respect to a text *T* [1..*n*] are an array MS[1..*m*] of tuples where MS[*i*].*len* is the length of the longest prefix *P* [*i*..*m*] that occurs in *T* and MS[*i*].*pos* is one location of that occurrence. Given MSs, *maximal exact matches* (MEMs) can be easily computed. Bannai et al. [6] showed how to compute MSs using an *r*-index by adding a *O*(*r*)-space *thresholds* structure encoding the positions of the minimum in the LCP array between two successive equal character runs. Given MS[*i*], their algorithm computes MS[*i*−1] by extending the match using LF if possible. On mismatch, thresholds and *lcp* queries are used to reposition to the “nearest” BWT run where LF can be used.

Rossi et al. showed how to find thresholds efficiently by using *prefix free parsing* [8] (PFP), applying them in MONI [23] alongside an *r*-index and a *straight-line program* (SLP) of size *g* supporting random access to *T* [11]. This approach computes MSs in *O*(*r* + *g*)-space and, given the insight of Baláž et al., *O*(*m* log *n*)-time with high probability. Ahmed et al.’s SPUMONI [2] introduced a derivative of MSs called *pseudo-matching lengths* (PMLs) using only the RLBWT and thresholds. The array PML[1..*m*] differs from MS lengths by always resetting the match length to 0 upon mismatch, thereby avoiding the use of SA samples and an SLP. PMLs were originally computed in *O*(*m* log log_𝒲_ (*n/r*)-time and *O*(*r*)-space before Zakeri et al.’s Movi [28] applied the move structure to improve the speed to *O*(*m*)-time. Whereas most queries require a BWT range, both PMLs and MSs track only a single BWT position and its corresponding run at every step.

### 2.5 Multi-Maximal Unique Matches

A *multi-maximal unique match* (multi-MUM) of a set of documents is a match occuring exactly once in each sequence that cannot be extended left or right without incurring some mismatch. This generalizes the pairwise notion of a MUM to *d >* 2 sequences. See Figure 1 for an example.

**Fig. 1.**
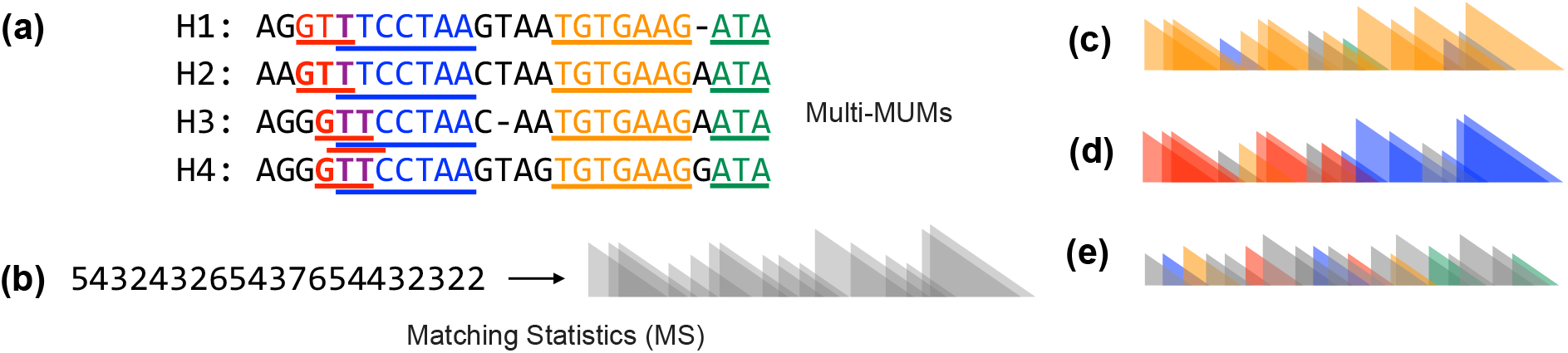
**(a)** A set of 4 sequences (“pangenome”) with multi-MUMs highlighted. Note that two multi-MUMs overlap. Also note that our method computes multi-MUMs from the sequences directly, and does not compute a multiple sequence alignment. **(b)** Matching statistic lengths (left) and their triangle-based schematic representation (right). **(c)** Schematic example of matching-statistics lengths derived from a query read and colored according to chain statistics. In this case, the chain statistics give a stronger basis for classifying the read as matching the reference index since matches are consistently from a single (orange) multi-MUM. **(d)** Similar example to (c) but for a read spanning adjacent (red and blue) multi-MUMs. **(e)** Example with no clear evidence of chaining, giving a stronger basis for classifying the read as not originating from the pangenome.

#### Definition 1.

*A multi-MUM between d >* 2 *sequences 𝒟* = {*T*_1_, *T*_2_, … , *T*_*d*_} *is defined by a width w and positions* [*p*_1_, *p*_2_, … , *p*_*d*_] *such that the set ℳ* = {*T*_*i*_[*p*_*i*_..*p*_*i*_ + *w* − 1] | ∀*i ∈* [1, *d*]} *are equal strings (i*.*e*., |M| = 1*) which are maximal:*

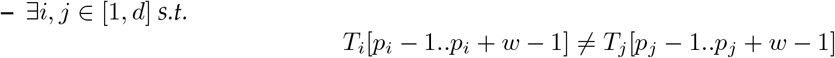

or

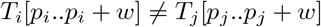

*and unique:*

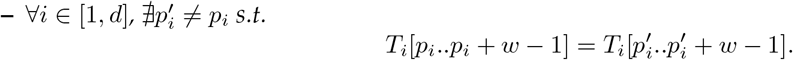

An alternative definition can be given in terms of SA ranges by assuming multi-MUM positions are with respect to a concatenated representation of the documents.

#### Theorem 1.

*Consider a collections of sequences* D = {*T*_1_, *T*_2_, … , *T*_*d*_} *and their concatenated text T* = *T*_1_*T*_2_ … *T*_*d*_$ *with* |*T* | = *n. A multi-MUM is present at* SA *position i ∈* [1, *n* − *d* + 1] *with w* = min{LCP[*i* + 1], … , LCP[*i* + *d* − 1]} *and positions* [*SA*[*i*], … , *SA*[*i* + *d* − 1]] *if and only if:*

1. *w >* LCP[*i*] *and w >* LCP[*i* + *d*]
2. |{DA[*i..i* + *d* − 1]}| = *d*
3. |{BWT[*i..i* + *d* − 1]}| *>* 1

The proof follows from Deogun et al. [10]. The *Mumemto* tool [25] finds multi-MUMs by verifying the conditions of Theorem 1, streaming windows of size *d* in *O*(*n*)-time and sublinear memory by using the same *prefix-free parsing* [8] method used by MONI to generate thresholds.

### 2.6 BWT Tunneling

Baier introduced the concept of *BWT tunneling* [3] as a compression technique for BWT encodings. It exploits *prefix intervals*: ranges of the BWT where characters visited through backwards stepping return identical substrings. We extend the framework to define the symmetric *suffix interval*.

#### Definition 2.

*A prefix interval [3] of* BWT[1..*n*] *is a position i with height h and width w such that*

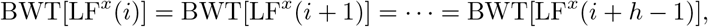

*for all* 0 ≤ *x < w*.

#### Definition 3.

*A suffix interval of* BWT[1..*n*] *is a position i with height h and width w such that*

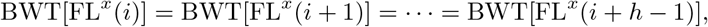

*for all* 0 ≤ *x < w*.

#### Corollary 1.

*If position i with height h and width w is a prefix interval, then position* LF^*w*−1^(*i*) *is a height-h, width-w suffix interval*.

Optimal tunneling involves choosing the subset of all possible prefix-intervals to maximize compression, a problem which is NP-hard if overlaps are permitted [4]. However, a non-overlapping set can be tunneled without a decision problem. We refer to a set of non-overlapping suffix intervals as *tunnels* due to this connection, but do not expand on the full procedure since our goal is not data compression.

#### Lemma 1.

*A set prefix intervals, represented by their corresponding suffix intervals, can be tunneled if no pair has overlapping* BWT *ranges such that when* (*i*_1_, *h*_1_, *w*_1_), (*i*_2_, *h*_2_, *w*_2_) *are distinct suffix intervals then*

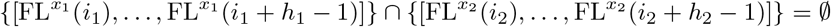

*for all* 0 ≤ *x*_1_ *< t*_1_, 0 ≤ *x*_2_ *< t*_2_.

*Proof*. From Baier’s definition of tunneling [3] and Corollary 1.

## 3 Methods

### 3.1 Properties of multi-MUMs

We can define the set of multi-MUMs by the criteria of Theorem 1. This allows us to bound the worst case number of multi-MUMs.

#### Definition 4.

*Let u be the number of multi-MUMs present within d concatenated sequences T* = *T*_1_*T*_2_ … *T*_*d*_$. *We can uniquely represent them by 𝒮*= [*s*_1_, … , *s*_*u*_] *and W* = [*w*_1_, … , *w*_*u*_], *where* ∀ *i∈* [1, *u*] *the range* SA[*s*_*i*_..*s*_*i*_+*d* − 1] *gives the starting positions of the multi-MUM in T and w*_*i*_ *gives its width*.

#### Lemma 2.

*Given the set of multi-MUMs as defined by Definition 4, all multi-MUM windows are non-overlapping such that for s*_*i*_, *s*_*j*_ *∈ 𝒮, if s*_*i*_ *< s*_*j*_ *then s*_*j*_ − *s*_*i*_ ≥ *d*.

*Proof*. Consider arbitrary *s*_*i*_ *∈ 𝒮, w*_*i*_ *∈ W* and assume for *k ∈* [1, *d* − 1] that *s*_*i*_ + *k ∈* S with corresponding *w*^*′*^. Then by Theorem 1 and *s*_*i*_ + *k* + 1 ≤ *s*_*i*_ + *d* ≤ *s*_*i*_ + *k* + *d* − 1,

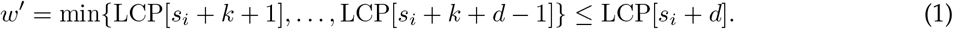

Similarly, *s*_*i*_ + 1 ≤ *s*_*i*_ + *k* ≤ *s*_*i*_ + *d* − 1 such that

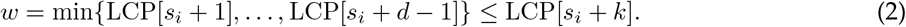

By Theorem 1 we have *w >* LCP[*s*_*i*_ + *d*] and therefore

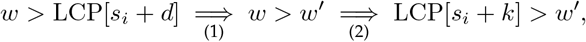

which contradicts Theorem 1, so *s*_*i*_ + *k ∈/* S. Hence, if *s*_*j*_ = *s*_*i*_ + *k ∈* S for *k >* 0 then *k* ≥ *d* and *s*_*j*_ − *s*_*i*_ = (*s*_*i*_ + *k*) − *s*_*i*_ = *k* ≥ *d*.

#### Corollary 2.

*For d sequences with* |*T* | = *n and u multi-MUMs, u* ≤ *n/d*.

#### Corollary 3.

*For a* BWT *of r runs with u multi-MUMs, u* ≤ *r*.

*Proof*. By Theorem 1 each multi-MUM with corresponding *s*_*i*_, *w*_*i*_ has |{*BWT* [*s*_*i*_..*s*_*i*_ + *w*_*i*_ − 1]}| *>* 1 implying at least two runs in the range. By Lemma 2 multi-MUM ranges cannot overlap, and hence *u* ≤ *r*.

### 3.2 Multi-MUM Tunnels

The SA range of each multi-MUM permits a maximum-width prefix interval with a corresponding suffix interval, found by forward stepping from the left end of the match.

#### Definition 5.

*For a multi-MUM with position s and width w let w*^*′*^ ≤ *w be the maximum value such that position* FL^*w′*^ (*i*) *is a height-d, width-w*^*′*^ *prefix interval. The multi-MUM-suffix interval for s, w is the height-d, width-w*^*′*^ *suffix interval*^3^ *at position* FL(*i*) *by Corollary 1*.

Although any single multi-MUM-suffix interval contains *d* · *w*^*′*^ BWT characters, there can exist overlaps between them. We can obtain a set of tunnels by removing all multi-MUMs which overlap; however, better coverage can be achieved by selectively truncating suffix interval widths^4^ upon collision. A set of tunnels induced from multi-MUM-suffix intervals permits only so many unique positions upon forward stepping, and we can find the unique set representing all multi-MUMs efficiently.

#### Definition 6.

*Let and W represent the multi-MUMs corresponding to a text with d sequences. Then positions 𝒮* _*t*_ = [*s*_1_..*s*_*u*_^*′*^ ] *and widths W*_*t*_ = [*w*_1_..*w*_*u*_^*′*^ ] *are a set of multi-MUM-tunnels if they satisfy Lemma 1 and they are induced from the set of multi-MUM-suffix intervals of Definition 5 such that:*

1. *The starting positions correspond to multi-MUM* SA *ranges, i*.*e*., {LF(*s*) | ∀*s∈*𝒮_*t*_}⊆{𝒮}
2. *Each tunnel has height d*.

#### Corollary 4.

*The set of starting positions found by forward stepping through all multi-MUM-tunnels is O*(*n/d*).

*Proof*. Definition 6 suggests non-overlapping intervals of size *d*. At most *n/d* of these can co-exist in a domain of size *n*.

#### Lemma 3.

*The unique set of multi-MUM-tunnels satisfying* {LF(*s*) | ∀*s∈*𝒮_*t*_}={𝒮} *is found in expected O*(*n/d* log log *n*)*-time and O*(*r* + *n/d*)*-space*.

*Proof*. By Lemma 2, no two multi-MUM SA ranges can overlap and marking the *O*(*n/d*) starting positions (Corollary 2) into a *y*-fast trie [27] with domain *n* takes expected *O*(*n/d* log log *n*)-time and *O*(*n/d*)-space. All multi-MUM-suffix intervals are forward stepped until completion if possible, or stopped if an *O*(log log *n*)-time predecessor query reveals an overlap . Corollary 4 ensures at most *O*(*n/d*) total FL steps in *O*(*n/d*)-time and *O*(*r*)-space using a move structure.

### 3.3 Co-linear BWT

We define the *co-linear BWT* (col-BWT)^5^ to be a division of the RLBWT which uses multi-MUM-tunnels to provide coarse-grained information. Let a sub-run refer either to a BWT run or a range contained within a BWT run. By Definition 2, every range in a tunnel corresponds to a sub-run of the BWT, allowing us to mark these ranges alongside existing runs (see Figure 2).

**Fig. 2.**
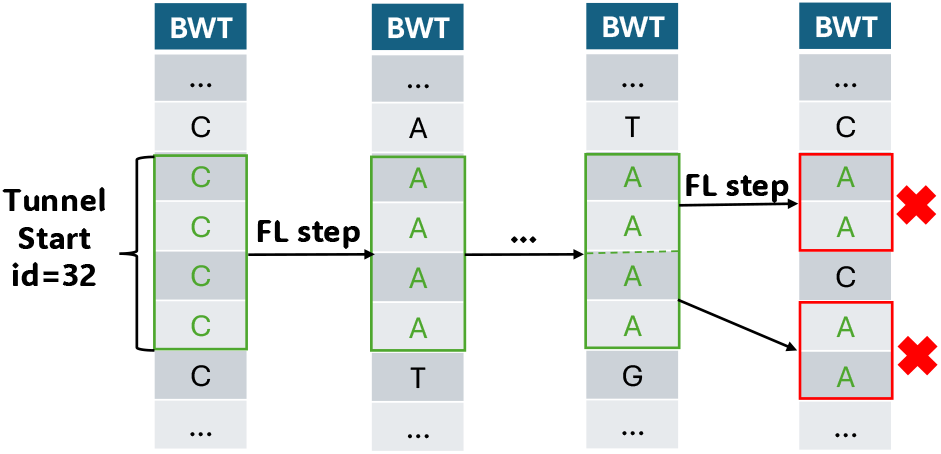
Forward stepping from multi-MUMs carves out sub-runs which can be marked with an *id*. We stop when FL steps are no longer contiguous, signifying the end of a tunnel, or truncated on overlap with another multi-MUM (not pictured). Tunnels were originally motivated as a strong predictor of BWT context [3], since backwards stepping anywhere within the range returns identical characters.

#### Definition 7.

*A co-linear* BWT *is an array* RLBWT_col_[1..*r*^*′*^] *of tuples representing all sub-run boundaries corresponding to:*

1. *Existing runs in* RLBWT[1..*r*].
2. *Ranges found by forward stepping through all multi-MUM-tunnels*.

*Let* RLBWT_col_[*i*].*c and* RLBWT_col_[*i*].*ℓ be the character and length of sub-runs respectively. We include an additional value* RLBWT_col_[*i*].*id which indicates which multi-MUM-tunnel the sub-run belongs to. If a sub-run does not correspond to a multi-MUM-tunnel it is given id 0*.

#### Theorem 2.

*Consider a* RLBWT_col_[1..*r*^*′*^] *built from a collection of d sequences where its* BWT *has r runs and length n. Then r*^*′*^ *∈ O*(*r* + *n/d*), *and further it can be represented in O*(*r* + *n/d*) *words of space*.

*Proof*. We initially have *r* runs. Let *P* be the set of distinct starting positions found by forward stepping through all multi-MUM tunnels, where |*P*|*∈ O*(*n/d*) by Corollary 4. For arbitrary *p∈ P* , either *p* corresponds to an existing run boundary or *p* marks a new sub-run: it introduces at most one new sub-run into the BWT. To mark the end of the sub-run, *p* + *d* similarly introduces at most one new sub-run. Since *p* is a tunnel (Lemma 1), ∄*i* with *p < i < p* + *d* such that *i* is the boundary of either a BWT run or another tunnel sub-run. It follows that the overall number of sub-runs inserted is at most *O*(*n/d*), and that *r*^*′*^*∈ O*(*r* + *n/d*). Corollary 3 allows log_2_ *r* bits per *id*, implying *O*(*r* + *n/d*) words of space.

### 3.4 Query Support

Recall that a move structure supports queries over permutations in space proportional to the number of contiguously permuted ranges; this still applies to sub-runs [20], which can be integrated into the move structure from a col-BWT to support LF in *O*(1)-time and *O*(*r* + *n/d*) words of space. Alternatively, an *r*-index can be adapted to use a col-BWT and compute LF in *O*(log log_𝒲_ *n/d*)-time and *O*(*r* + *n/d*)-space. Sub-runs fit into PML or MS computation without issue: threshold jumping is not hindered [28], and BWT positions can be framed with respect to sub-runs to find an *id* if present. This result matches the query speeds of Section 2.4 while outputting coarse-grained co-linearity statistics which we call *chain statistics*.

#### Corollary 5.

*Given a pattern P* [1..*m*], *a col-*BWT *can be used to output chain statistics* CID[1..*m*] *in O*(*m*)*-time and O*(*r* + *n/d*)*-space, where* CID[*i*] *is the multi-MUM-tunnel that a corresponding* MS[*i*].*pos occurs in. If using* MSs *to find MEMs, we can identify which multi-MUM-tunnel its corresponding* MS *occurrence ends in. It supports computing* PML*/*MS *in the same complexity as previous best approaches (Section 2.4) by replacing an O*(*r*)*-space index with an O*(*r* + *n/d*) *representation*.

### 3.5 Implementation

Our algorithm to identify sub-runs follows closely from Lemma 3. Mumemto [25] is used to generate the SA ranges and widths of multi-MUMs for a collection of sequences; since it uses PFP, we also output the RLBWT and thresholds simultaneously. The multi-MUM SA ranges are then marked in a bitvector supporting predecessor queries to identify overlaps. To support forward stepping, a move structure over FL is built; this is fast in practice and theory since by definition tunnels FL step as a range. We find tunnels by forward stepping from every multi-MUM SA range until we reach an overlap or the end of the multi-MUM-suffix interval. This process inserts sub-runs alongside multi-MUM-tunnel *id*s, which are used to build a modified version of Movi [28]. We also support finding all sub-runs corresponding to multi-MUMs by relaxing the tunnel constraint, although this is worst case *O*(*n*)-space. The repository is available at https://github.com/drnatebrown/col-bwt.

Using log_2_ *r*-bits per *id* might be wasteful; to identify if matches identified through fine-grained queries belong to the same multi-MUM it suffices to check if their *id*s are equal. Using fewer bits permits false positives, but not false negatives. Our index bins *id*s into one byte, since this was not found to adversely affect the application of chain statistics (appendix, Figure A1). Further, since all sub-runs corresponding to tunnels have trivial height *d*, we can re-use the space Movi allocates to run lengths and replace it with an *id* when necessary. A further space reduction is achieved by sub-sampling. Consider marking a sub-run every *s* steps from the start of a tunnel, thereby reducing the overall number of sub-runs needed; this still ensures at least one sampled sub-run is hit if a match is extended with ≥ *s* consecutive LF steps.

## 4 Results

The results of Theorem 2 and Corollary 5 describe an index that is *O*(*r* + *n/d*) words of space. However, the ability to output meaningful chain statistics is dependent on the multi-MUM coverage of a dataset. To evaluate, we implemented the procedure and performed a series of experiments using haplotypes from the year-1 freeze of the Human Pangenome Reference Consortium [16] (HPRC). We tested multi-MUM coverage in practice by “splitting” runs for up to 94 haplotypes. Each genome and its reverse compliment are considered a single document. Figure 3 shows the overall percentage of sequences which are given a multi-MUM *id* by our index and the number of resulting sub-runs after splitting^6^.

**Fig. 3.**
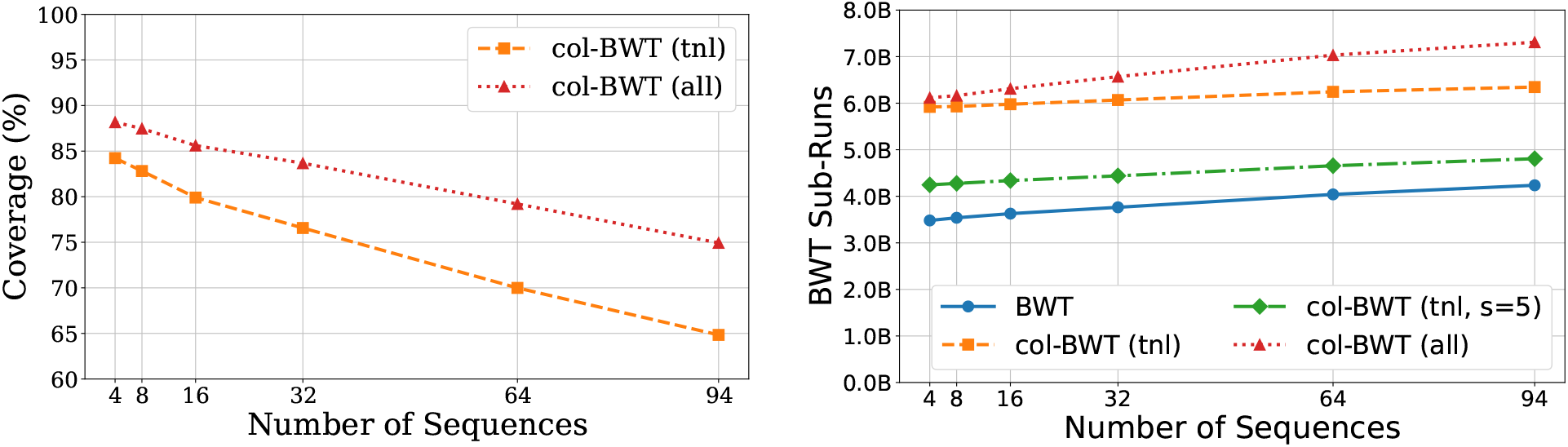
**Left:** Percentage of characters covered by a sub-run when considering only multi-MUM-tunnels (`tnl`) or all multi-MUM sub-runs (`all`) using HPRC collections. **Right:** The number of sub-runs (or runs, for `BWT`) in billions resulting from marking multi-MUMs of HPRC collections, where *s* is the sub-sampling parameter.

### 4.1 Applying Chain Statistics

We tested whether chain statistics can correctly determine if two adjacent matches in the fine-grained statistics (PMLs in this case) could be classified as co-linear or not. Since a PML match is either extended by 1 or dropped to 0, they form a number of “peaks”^7^. When a mismatch occurs due to error or slight variation in the read, we may observe small peaks until reoriented in the BWT. This scenario can produce adjacent “significant” peaks (i.e., of meaningful length) which are co-linear with respect to the reference but not captured using solely fine-grained statistics. Our approach is to classify adjacent peaks, above some threshold to ignore small matches, as co-linear if the *id* at the end of the first peak matches the *id* at the start of the next. This approach works well with sub-sampling: if parameter *s* is less than or equal to the peak threshold value we are guaranteed to identify the necessary *id* to classify.

To evaluate, we indexed the chromosome 19 components of HPRC and simulated 10,000 reads of length 10kbp from the reference with SNV rate 1%. Figure 4 shows the resulting accuracy, where co-linear peaks separated by SNVs are true positives. We find accuracy correlates with coverage when marking all multi-MUM sub-runs and over performs with respect to coverage for multi-MUM-tunnels.

**Fig. 4.**
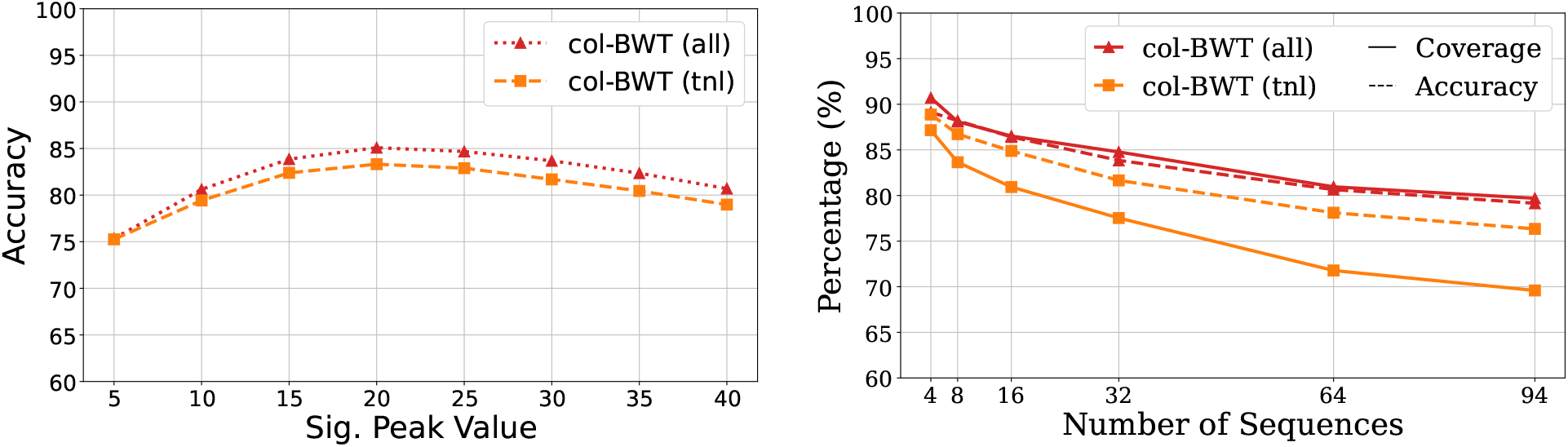
Results for splitting when considering only multi-MUM-tunnels (`tnl`) or all multi-MUM sub-runs (`all`). **Left:** The accuracy of identifying co-linear peaks using 32 HPRC chromosome 19 copies using 10kbp reads with SNV rate of 1%. **Right:** The accuracy across collections of HPRC chromosome 19 copies alongside coverage of *id* sub-runs using a significant peak value of 20.

### 4.2 Read Classification

#### Classification Scheme

We hypothesized earlier that by considering chain statistics in combination with finer-grained statistics such as MSs or PMLs, we could improve our classification accuracy to be comparable to that of alignment-based methods that perform explicit chaining. Ahmed et al’s *SPUMONI 2* [1] uses PMLs to classify reads by first constructing a *null PML distribution*. Substrings from the reference are extracted, reversed, and then used to generate PMLs back against the reference. These serve as random sequences (they are not the reverse compliment) that share the same distribution of bases as the reference. Ahmed et al. [1] consider the largest PML occurring at least 5 times in the null distribution to be a threshold PML value *k*. Reads are divided into non-overlapping windows, by default 150bp, and classified as matching to the reference if the majority of windows contain a PML greater than *k*.

The peak identification scheme of Section 4.1 can be used alongside PMLs to perform classification. Chain statistics are generated for the null sequences, and a significant peak threshold is used to find the expected number of peaks classified as co-linear (ratio-ed by read length). The significant peak threshold is set to ⌈(*k* + 1)*/*2⌉by default: intuitively, this attempts to identify peaks whose cumulative height must be greater than *k*. We consider three formulations which combine these methods, classifying a read to the reference if: **a)** the number of peaks classified as co-linear is greater than the null expected; **b)** the majority of windows after augmenting PMLs through “merging” the height of adjacent co-linear peaks contain a PML greater than *k*; **c)** either the majority of windows contain a PML greater than *k*, or the the number of peaks classified as co-linear is greater than the null expected.

The original SPUMONI 2 scheme can be used for *adaptive sampling* [1], since its classification method can be applied while the read is streamed in and computation paused/resumed at any time. The methods we have introduced, used concurrently with PMLs, inherit the same advantages.

#### Experiments

We replicated the experiment of Ahmed et al.’s original SPUMONI 2 study [1]. 500,000 simulated human reads are generated from the CHM13 reference [21] using PBSIM3 [22] with mean accuracy 95% and combined with 500,000 real nanopore reads from human gut microbial species [18]. This models *host depletion* where the goal is to remove human host reads and retain microbial reads. Our result does not model adaptive sampling, differing from the original study by providing full reads rather than batching. We consider successful classification of human reads to be true positives.

The following indexes were built on HPRC collections of 10 and 32 haplotypes: `SPUMONI`, the SPUMONI 2 index [1] without minimizer digestion^8^ and *k* = 5; the default `Movi` [28] construction supporting PML computation with prefetching; `minimap2` [15] built using ONT read presets and three threads; `col-BWT` as outlined in Section 3.5 using multi-MUM-tunnels, *s* = 10, *k* = 5, and otherwise the same settings as `Movi`. The time and maximum memory were measured using GNU time on a server with an Intel(R) Xeon(R) Gold 6248R CPU running at 3.00 GHz with 48 cores and 1.5TB DDR4 memory. Table 1 shows the resulting metrics; the best classification approach of `col-BWT` improves over fine-grained statistics to reach the same percentile accuracy as `minimap2`.

**Table 1.** Read classification metrics when modeling host sequence depletion using HPRC collections of 10 (top) and 32 (bottom) haplotypes with reverse compliment. `Movi` uses the same classification approach as `SPUMONI`, so some measurements are omitted (shown as “–”); similarly, `minimap2` versions differ only by maximum memory usage. Multiple classification schemes are presented for `col-BWT`, corresponding to those labeled in Section 4.2. The `SPUMONI` threshold PML value was found to be 19 in both datasets, resulting in a significant peak threshold of 10 for `col-BWT` approaches. Index size is measured as its total disk footprint. See appendix, Table A2, for using all multi-MUM subruns; improvements in accuracy are slight in comparison with increased time/memory usage.

| HPRC 10 | SPUMONI | Movi | col-BWT<sup>a</sup> | col-BWT<sup>b</sup> | col-BWT<sup>c</sup> | minimap2 | minimap2<sup>†</sup> |
| --- | --- | --- | --- | --- | --- | --- | --- |
| Accuracy | 96.54 | - | 96.16 | 98.75 | 99.69 | 99.96 | - |
| Sensitivity | 93.08 | - | 92.33 | 97.52 | 99.41 | 99.99 | - |
| Specificity | 99.98 | - | 99.97 | 99.98 | 99.98 | 99.93 | - |
| Index Size (GB) | 10.08 | 41.83 | 46.30 | - | - | 130.11 | 119.62 |
| Peak Mem. (GB) | 10.82 | 41.84 | 46.31 | - | - | 20.51 | 122.69 |
| Time (minutes) | 622.36 | 23.97 | 22.97 | - | - | 920.28 | 662.89 |

| HPRC 32 | SPUMONI | Movi | col-BWT<sup>a</sup> | col-BWT<sup>b</sup> | col-BWT<sup>c</sup> | minimap2 | minimap2<sup>†</sup> |
| --- | --- | --- | --- | --- | --- | --- | --- |
| Accuracy | 96.55 | - | 95.12 | 98.56 | 99.25 | 99.96 | - |
| Sensitivity | 93.11 | - | 90.25 | 97.14 | 98.52 | 99.99 | - |
| Specificity | 99.98 | - | 99.98 | 99.98 | 99.98 | 99.92 | - |
| Index Size (GB) | 12.58 | 44.10 | 48.27 | - | - | 413.66 | 378.26 |
| Peak Mem. (GB) | 13.32 | 44.11 | 48.28 | - | - | 21.54 | 383.98 |
| Time (minutes) | 490.26 | 13.30 | 17.17 | - | - | 3273.94 | 2194.16 |

<sup>†</sup> Loads full index into memory, instead of default 8GB split indexes.

## 5 Discussion

We presented the first framework and method for simultaneous computation of fine-grained matching statistics (or pseudo-matching lengths) and coarse-grained chain statistics. We proved that the addition of the chain statistics can be done while achieving a space bound of *O*(*r* + *n/d*), where *r* grows with the amount of distinct sequence in the collection and *n/d* is the average length of a sequence in the collection. It was shown previously that *r* typically grows much more slowly than *n* as the number of similar sequences in the collection grows. Further, *n/d* stays approximately constant as the collection grows.

We showed that computing PMLs and chain statistics simultaneously achieves the same linear-time bound as computing PMLs alone. Remarkably, this allows for distinct matches to be “chained” (i.e. found to either be co-linear or not co-linear with respect to the collection) in linear time, which contrasts with typical chaining approaches that require *O*(*n* log *n*) time with respect to the number of seeds *n*, or worse. Finally, we showed that the addition of chain statistics allows for more accurate classification of reads, ultimately achieving an accuracy that is comparable to that of a slower alignment method, minimap2 [15]. A drawback of our approach is that the ability to cover a genome with multi-MUMs decreases as the number of indexed genomes increases. That is, adding additional genomes tends to fragment or eliminate multi-MUMs such that they cover a lower proportion of the overall sequence. This is visible in the differences between our results for 10 HPRC haplotypes versus those for 32 HPRC haplotypes. This points to the need to include one of two extensions to our method. First, we could let the marking of the multi-MUMs “smear” to surrounding areas that are not strictly part of a multi-MUM but are near to one. Second, we could perform a priori chaining of multi-MUMs to cover polymorphic regions between them, similarly to how a multiple sequence aligner would chain matches to form longer approximate matches. By doing this, we can shift from operating strictly at the level of multi-MUMs to a coarser level of approximate chains-of-multi-MUMs, which we expect to cover a larger proportion of the bases of the pangenome.

Further, requiring non-overlapping multi-MUMs is not proven to be a strict requirement. Although multi-MUM-tunnels are well motivated in reference to past work, it is unclear if it is required to remove overlaps to achieve a sub-linear space bound. Schemes which do not omit overlaps will lead to higher coverage, which may further improve applications. We could also consider relaxing the strictness imposed by multi-MUMs to instead find maximal exact matches that occur more than once in a given document, or in only some *d*− *k* documents. Both methods should improve coverage in practice. Considering subsets of documents may also be preferred when some sequences have preferred traits to identify. This also applies to modular indexes, where multi-MUMs are found for subsets of the collection instead and then queried. Exploration of efficient techniques to merge multi-MUMs would further enrich these methods, while also supporting incremental additions of documents into the index.

Our method is related to Wheeler maps [5], which associate arbitrary “tags” with positions in a compressed BWT index. However, we avoid the overhead introduced by the fact that tag runs will generally fail to coincide with BWT runs. Wheeler maps necessitate additional data structures and queries to relate to tagged run boundaries; in comparison, our splitting procedure modifies the BWT runs in a way that forces them to coincide precisely with multi-MUM labels, which allows linear-time queries. However, Wheeler maps provide a bound on document listing. In theory, the methods could be combined to use multi-MUM sub-runs as tags while achieving a sub-linear space bound, something Wheeler maps do not inherently possess (see appendix section A1).

Of independent interest is our ability to generate sets of non-overlapping tunnels. To avoid the hardness of finding an optimal tunneling, previous heuristics [3] avoid overlaps by generating tunnels whose starts and ends coincide exactly with the range of a BWT run. Our method can also be seen as a heuristic for finding BWT tunnels, though ours is less restrictive in that it allows tunnels that may “carve out” a sub-run from within an original run. Compressing multi-MUM tunnels corresponds to collapsing identical stretches shared by all documents, which is encouraging given the coverage results observed by our method. However, coverage alone is not a strong enough indicator that BWT tunneling performs well, meaning evaluation of our approach when applied to compression is required.

Finally, we emphasize that our method does not use multiple sequence alignment (MSA). Rather, multi-MUMs can be computed as a by-product of computing the BWT of the input sequence collection. Thus, col-BWT avoids the drawbacks associated with methods that first build an MSA, which can require removal of substrings that are overly repetitive or overly inconsistent with a global coordinate system.

## Acknowledgments

N.K.B. was supported by a Johns Hopkins University Computer Science PhD Fellowship. N.K.B. and B.L. were supported by NIH grants R21HG013433 and R01HG011392 to B.L. V.S. was supported by the National Science Foundation grant DGE2139757. We thank Mohsen Zakeri and Omar Y. Ahmed for helpful technical discussions.

**Appendix**

### A1 Wheeler Maps

#### Lemma 4.

*The sub-runs of the* RLBWT_col_ *given in Definition 7 can be used to build a Wheeler map [5] which, for a pattern P* [1..*m*] *after O*(*m* log *n*) *time preprocessing, can list the k distinct multi-MUM-tunnel ids for for any P* [*i..j*] *in optimal O*(*k*)*-time in O*(*r* + *g* + *n/d*)*-space, where r is the number of runs in the* BWT *and g the size of an SLP built over the text*.

*Proof*. A Wheeler map supports the above operations in *O*(*r* + *g* + *t*)-space where *t* is the number of runs in an arbitrary array tagging characters of the BWT [5]. Substituting an RLBWT_col_, containing *O*(*r* + *n/d*)-runs, allows us to do so in *O*(*r* + *g* + *n/d*)-space where tags are multi-MUM-tunnel ids.

#### Lemma 5.

*Given a pattern P* [1..*m*], *a tag array of t runs, as defined for a Wheeler map [5], can be used to output respective tags in O*(*m*)*-time where a tag corresponds to the occurrence of* MS[*i*].*pos. If using* MSs *to find MEMs, we can identify which tag its corresponding* MS *occurrence ends in. It supports the operation in O*(*r* + *t*)*-space*.

*Proof*. This abstracts Corollary 5 for an arbitrary *t*. Here the number of runs it contains cannot necessarily be bound, nor do tag runs necessarily coincide with BWT runs in a way to provide a bound.

### A2 Tables

**Table A1.** Elapsed time to identify multi-MUM tunnels and output corresponding sub-runs and *id*s to file, measured in minutes. Timings refer to the experiment of Figure 3.

| # HPRC Seqs: | 4 | 8 | 16 | 32 | 64 | 94 |
| --- | --- | --- | --- | --- | --- | --- |
| tnl | 71.79 | 98.72 | 80.03 | 89.65 | 109.34 | 90.19 |
| tnl, (s=5) | 49.86 | 67.93 | 50.40 | 54.20 | 60.86 | 62.89 |
| all | 86.12 | 103.16 | 77.95 | 112.16 | 110.56 | 120.46 |

**Table A2.** The same experiments and results as Table 1 for 32 HPRC haplotypes, except using a `col-BWT` with all multi-MUM sub-runs and *s* = 10. This method obtains slightly higher metrics than the multi-MUM tunnel approach shown in the main table, likely due to higher coverage. However, it suffers in time/memory usage.

| HPRC 32 | SPUMONI | Movi | col-BWT<sup>a</sup> | col-BWT<sup>b</sup> | col-BWT<sup>c</sup> | minimap2 | minimap2<sup>†</sup> |
| --- | --- | --- | --- | --- | --- | --- | --- |
| Accuracy | 96.55 | - | 95.35 | 98.62 | 99.27 | 99.96 | - |
| Sensitivity | 93.11 | - | 90.72 | 97.26 | 98.58 | 99.99 | - |
| Specificity | 99.98 | - | 99.97 | 99.98 | 99.98 | 99.92 | - |
| Index Size (GB) | 12.58 | 44.10 | 66.32 | - | - | 413.66 | 378.26 |
| Peak Mem. (GB) | 13.32 | 44.11 | 66.32 | - | - | 21.54 | 383.98 |
| Time (minutes) | 490.26 | 13.30 | 21.17 | - | - | 3273.94 | 2194.16 |

**Fig. A1.**
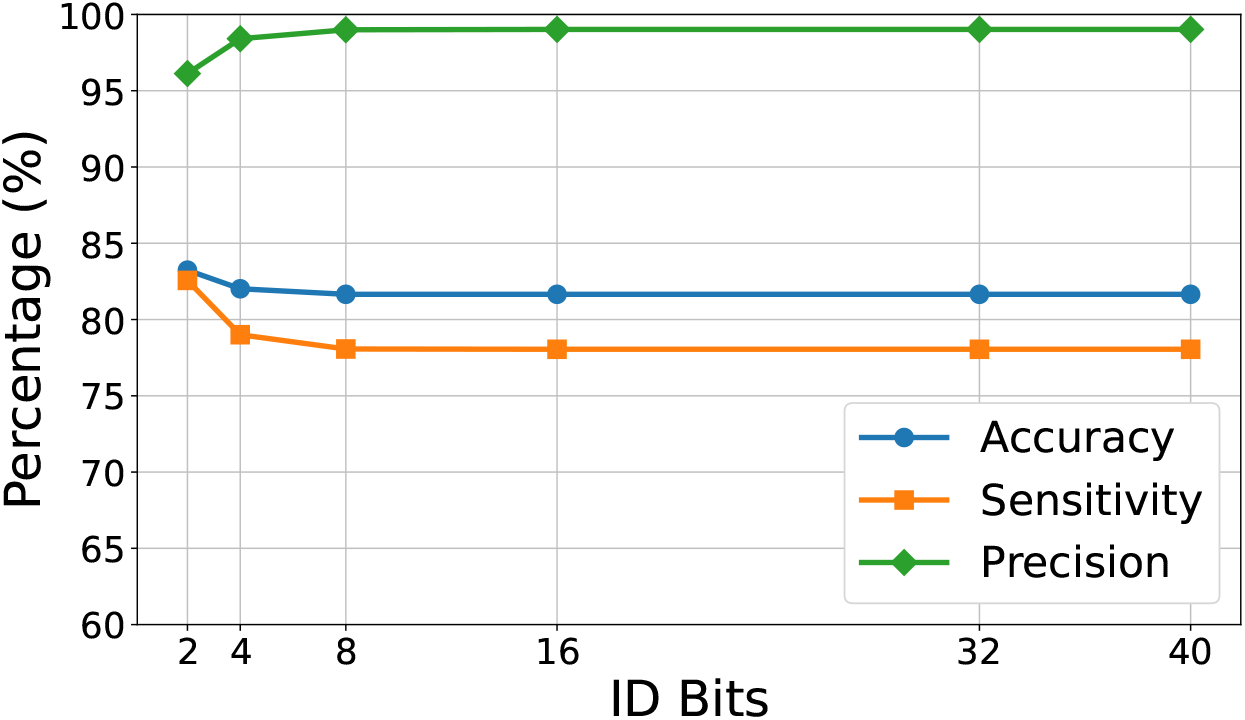
Explores the experiments of Section 4.1 by varying the number of bits allocated to store *id*’s. The metrics are with respect to classifying co-linear peaks of 32 HPRC chromosome 19’s using 10kbp reads with SNV rate of 1% and significant peak value 20.

## Footnotes

1 Separators between documents ignored for convenience, see [7] for a full study on multi-string BWTs.

2 For polylogarithmic alphabets where *σ* = *O*(polylog *n*).

3 The multi-MUM SA range and BWT range are off by one FL step; by definition BWT precedes SA characters.

4 Truncating heights leads to higher coverage, but loses the connection to multi-MUM SA ranges.

5 We use *col* as it abbreviates co-linearity (not column). A *col* is also the lowest point on a ridge between two peaks, which describes our index aptly. In construction, we carve runs from a larger range due to their heightened similarity. In application, we bridge gaps between peaks when they appear to be of the same landscape (Figure 1).

6 The running times are provided in appendix, Table A1.

7 MSs also contain peaks with a decrease afterwards; however, the value after the peak is not necessarily 0.

8 Movi does not support minimizer alphabets and we omit this option for comparison. Since these improve sensitivity by reducing precision, improvements in accuracy by our methods should also be observed in minimizer digests.

